# Structural basis for recognition of diverse antidepressants by the human serotonin transporter

**DOI:** 10.1101/204859

**Authors:** Jonathan A. Coleman, Eric Gouaux

## Abstract

Selective serotonin reuptake inhibitors are clinically prescribed antidepressants that act by increasing the local concentration of neurotransmitter at synapses and in extracellular spaces via blockade of the serotonin transporter. Here we report x-ray structures of engineered thermostable variants of the human serotonin transporter bound to the antidepressants sertraline, fluvoxamine, and paroxetine. The drugs prevent serotonin binding by occupying the central substrate binding site and stabilizing the transporter in an outward-open conformation. These structures explain how residues within the central site orchestrate binding of chemically diverse inhibitors and mediate transporter-drug selectivity.

## MAIN TEXT

In the central nervous system the serotonin transporter (SERT) modulates serotoninergic signaling by carrying out the uptake of serotonin (5-HT) from the synaptic space into presynaptic neurons^1^. 5-HT influences many aspects of behavior including memory, learning, sleep, hunger, pain, sexual function, and mood^2^. 5-HT is also found in the circulatory system where it acts as a vasoconstrictor, and altered plasma concentrations of 5-HT have been implicated in several pathologies, including hypertension^3^. The plasma levels of 5-HT are regulated by SERT, which stores 5-HT in platelets, ensuring a stable blood flow^4^. Transporters for dopamine (DAT) and norepinephrine (NET) are related to SERT in amino acid sequence and in biological function and are also well-established targets for pharmacological agents that influence brain chemistry^5^. These monoamine transporters exploit the ion gradients of Na^+^ and Cl^−^ in order to render neurotransmitter (NT) uptake thermodynamically favorable^6^. Monoamine transporters have twelve transmembrane helices (TM) with TMs 1–5 and TMs 6–10 forming an inverted-topological repeat within which is a central binding site for substrate and ions approximately halfway across the membrane^7–9^. An extracellular vestibule in the outward-facing conformation^7,9,10^ forms a secondary, allosteric site which, when occupied by ligands, can result in modulation of transporter activity by altering the kinetics of ligand dissociation from the central site^11–13^. Addictive substances such as cocaine and amphetamine bind to monoamine transporters and can either inhibit NT transport or promote NT efflux, respectively^14,15^. Selective serotonin reuptake inhibitors (SSRIs) are a class of small molecules that are highly selective for SERT over DAT and NET and that inhibit 5-HT reuptake with nanomolar potency^16^. SSRIs are typically used as antidepressants in the treatment of major depressive disorder and anxiety disorder^1^ and are the most widely prescribed antidepressants^17^.

SSRIs have a diverse chemical structure and, in many instances, they do not share common structural motifs. The diversity of SSRI chemical structures, in turn, results in compounds with substantial pharmacological differences^18^. Sertraline and fluvoxamine, as examples, differ in chemical structure from other SSRIs such as paroxetine, citalopram, and fluoxetine (Fig. 1), and as a consequence these compounds bind with a range of affinities to SERT. Sertraline contains a tetrahydronaphthalene ring system linked to a secondary amine together with a meta and para substituted dichlorophenyl group. Fluvoxamine, by contrast, consists of a 2-aminoethyloxime moiety attached to a methyoxybutyl group, and a phenyl group containing a trifluoronated methyl at the *para* positon. Fluoxetine also contains a trifluoronated aromatic group but is instead coupled to a phenylpropylamine moiety. Paroxetine and citalopram also differ substantially in the structures of their aromatic, amine, and halogenated substitutents.

**Figure 1.**
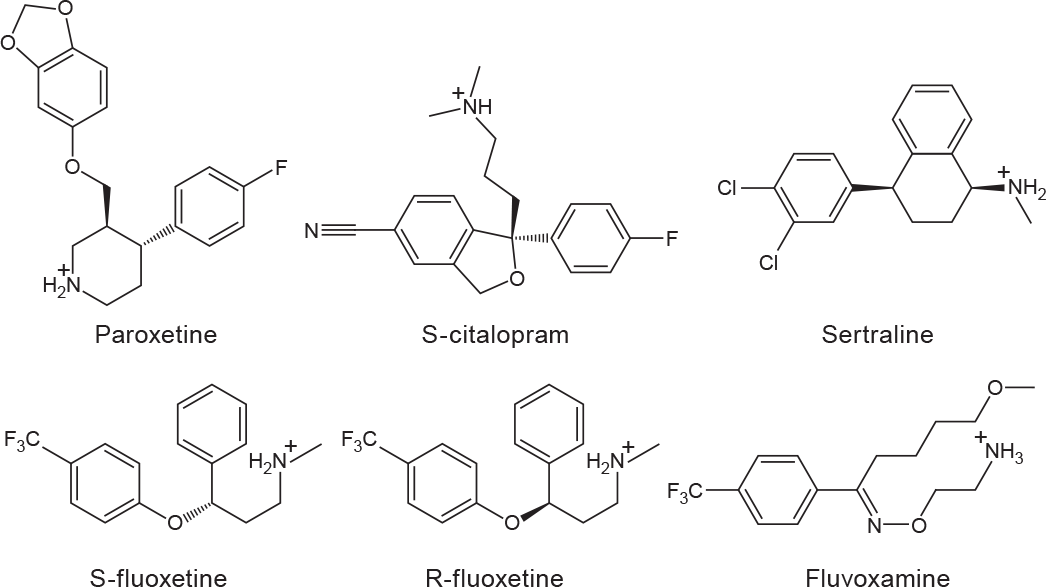
Chemical structures of SSRIs.

Recently, we solved x-ray structures of a thermostabilized, transport-inactive construct of human SERT, deemed the ts3 construct, in complex with the SSRIs paroxetine and citalopram^8^. We employed the thermostabilized variant of SERT in order to facilitate purification and crystallization^19,20^. One of the thermostabilizing mutations, however, involves a residue (Thr439Ser) that is directly positioned within the central binding site, in close proximity to the bound SSRIs. Moreover, recent computational modeling of a wild-type SERT-paroxetine complex, in which the structure of the *Drosophila* DAT was used as a template for the SERT structure, yielded a pose for paroxetine in the SERT binding site that is different from that found in the SERT ts3 crystal structure ^21^. In light of the computational study and because residue 439 is near the central binding site, we were motivated to determine crystal structures of SERT bound with SSRIs where residue 439 is the wild-type threonine amino acid.

Despite the importance of SSRIs in medicine and in the biophysical study of SERT, there is little structural explanation for how these diverse ligands bind to SERT ^22^. Until recently, the structural understanding of how SSRIs bind to SERT has been largely guided by studies of the bacterial homolog LeuT^23,24^, computational modeling^25–29^, and more recently, the structures of SERT in complex with paroxetine and citalopram^8^. In order to accommodate various SSRIs within the central binding site, we reasoned that residues lining the central binding site might adopt different conformations. Thus we determined x-ray structures of Thr439 ts2 SERT in complex with paroxetine and the structure of the ts3 transporter bound to sertraline and fluvoxamine. These structures provide insight into how different SSRIs are bound within the central binding site of SERT.

To probe the capacity of the ts3 and ts2 transporters to bind antidepressants we first carried out binding studies using ^3^H-paroxetine (Fig. 2a). Paroxetine binds with a dissociation constant (K_d_) value of 1.09 ± 0.08 and 0.63 ± 0.07 nM to the ts3 and ts2 variants (student’s t-test, p-value less than 0.05). Next, we measured the K_i_ values for various SSRIs by competition with ^3^H-paroxetine. With the exception of sertraline, SSRIs bind with slightly lower affinity to ts3 *vs.* ts2 (Fig. 2b). S-citalopram exhibited a K_i_ of 10 ± 1 and 7 ± 1 nM (p-value greater than 0.05); sertraline, 2.0 ± 0.2 and 7 ± 1 nM (p-value greater than 0.05); S-fluoxetine, 35 ± 3 and 10 ± 1 nM (p-value less than 0.05); R-fluoxetine, 41 ± 4 and 9 ± 1 nM (p-value less than 0.05); and fluvoxamine, 69 ± 9 and 10 ± 2 nM (p-value less than 0.05). These dissociation constants and the relative differences of the dissociation constants between the SSRIs agree well with previous reports^30^. The presence of the Y110A mutant in both the ts3 and the ts2 variants renders them inactive in 5-HT transport^19^.

**Figure 2.**
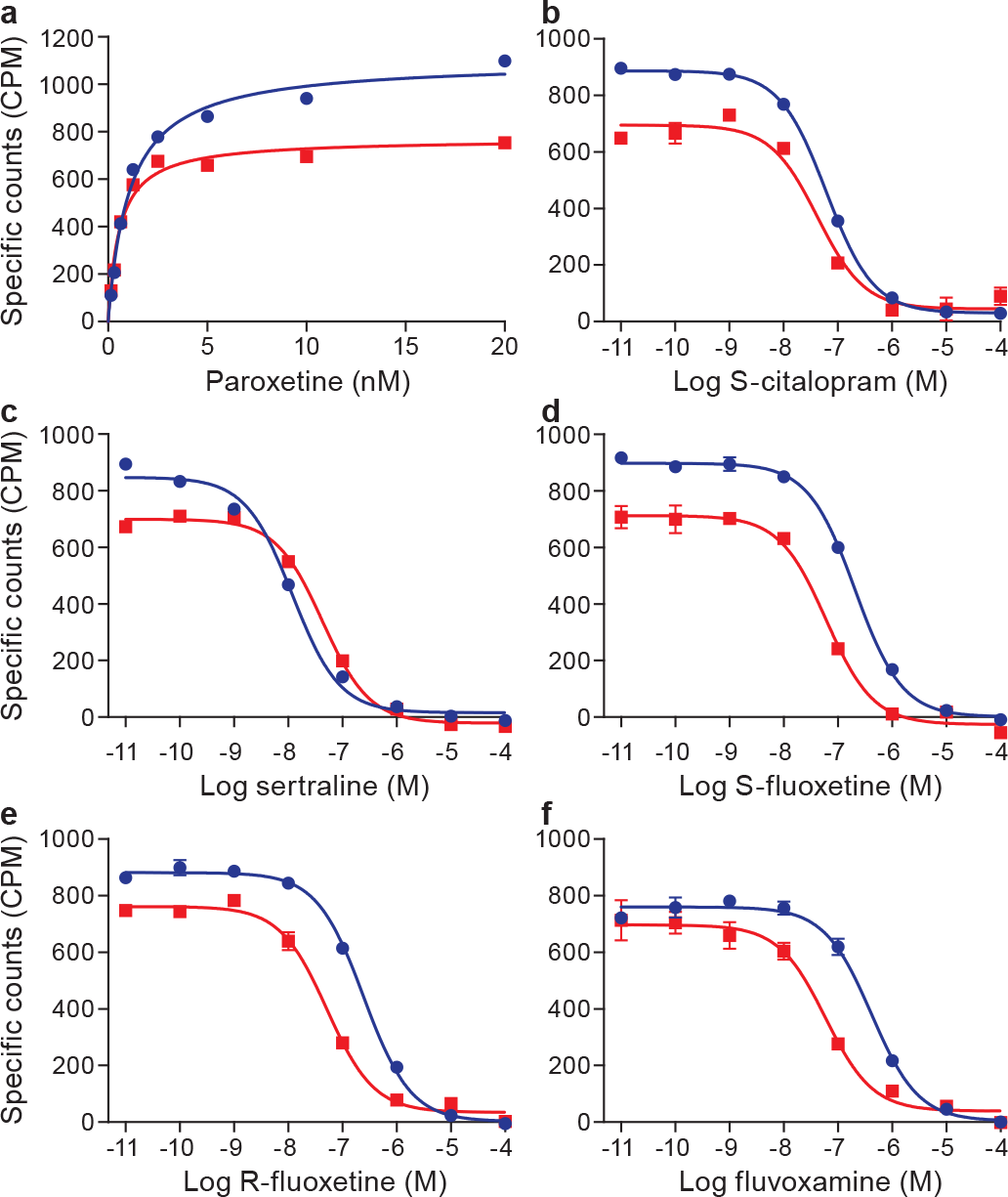
Saturation and competition binding experiments. In **panel a** are shown the saturation binding of ^3^H-paroxetine to ts3 (red, squares) and to ts2 (blue, circles) variants. **Panels b-f** show competition experiments between the binding of ^3^H-paroxetine and cold SSRIs, also on the ts3 and ts2 SERT variants, as described in **panel a.** Graphs depict the average of triplicate measurements from a representative experiment (error bars represent s.e.m.).

The structure of paroxetine bound to ts2 was determined using diffraction data that extend to Bragg spacing of 3.6 Å. Electron density for several of the side chains for residues in the intracellular gate and within the C-terminal hinge and helix were also better resolved in the ts2 structure reported here in comparison to the previously reported ts3 structure, thus allowing for more complete modeling of these regions (Extended Data Fig. 1), including part of the recognition sequence for SEC24C^31,32^. Despite the medium resolution of the diffraction data, the electron density features in the central site are of sufficient quality to allow us to position paroxetine. We find that the piperidine ring is best accommodated in subsite A, the benzodioxol in subsite B, and the fluorophenyl group in subsite C (Fig. 3a, Table 1). Fitting of paroxetine in the opposite orientation with the benzodioxol in subsite C, and the fluorophenyl in subsite B yields a poor fit to the electron density and produces clashes in subsite A and C (Extended Data Fig. 2). The sertraline and fluvoxamine structures were solved in complex with ts3 at 3.5 and 3.8 Å respectively. Omit maps and anomalous difference Fourier maps were used to position sertraline with the dichloro phenyl ring in subsite B and the amine and tetrahydronaphthalene groups in subsites A and C, respectively (Fig. 3b). The density for fluvoxamine was not continuous but we found density for the aminoethyloxime and trifluoro aromatic groups in subsites A and B (Fig. 3c); no density was observed for the methyoxybutyl moiety, likely due to its flexibility. A Polder ‘omit’ map^33^ was used to further recover fluvoxamine density and position it more accurately within subsites A and B. Placement of residues which are involved in drug binding at the central site was guided by electron density features and by chemical interactions with the drug (Extended Data Fig. 3). No density was observed in the allosteric site with the sertraline and fluvoxamine complexes, consistent with sertraline being a less potent allosteric inhibitor of the dissociation of citalopram from the central site compared to citalopram^12^. At present there is no information on whether fluvoxamine binds to the allosteric site and influences ligand dissociation from the central site.

**Figure 3.**
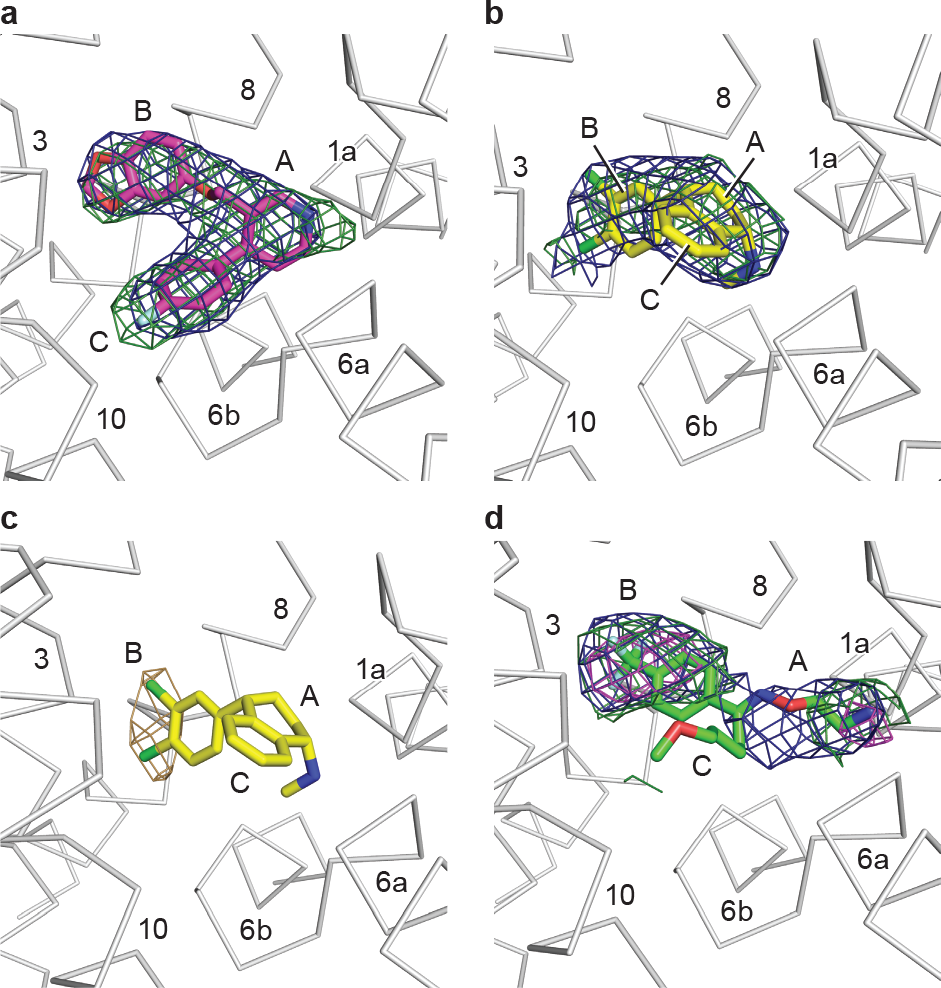
Antidepressant binding in the central binding site. **a**, Shown is the ‘omit’ electron density (green mesh) associated with paroxetine, contoured at 4σ. Electron density derived from a Polder ‘omit’ electron density is shown in blue mesh and is contoured at 7σ. **b**, This shows electron density for sertraline derived from an ‘omit’ map, contoured at 2.5σ; the Polder ‘omit’ electron density is contoured at 5σ. **c**, Anomalous difference electron density (light brown), derived from sertraline, is contoured at 3σ. **d**, Density for fluvoxamine derived from an omit electron density map and contoured at 1σ (green) and 2σ (magenta), with the Polder omit electron density contoured at 3σ. The approximate positions of subsites A, B, and C are shown.

**Table 1.** Data collection and refinement statistics.

|  | Paroxetine S439T<br>6AWN | Fluvoxamine ts3<br>6AWP | Sertraline ts3<br>6AWO | Sertraline ts3<br>6AWQ |
| --- | --- | --- | --- | --- |
| **Data collection** | APS24-IDE | APS24-IDE | APS24-IDE | ALS 5.0.2 |
| Space group | C222<sub>1</sub> | C222<sub>1</sub> | C222<sub>1</sub> | C222<sub>1</sub> |
| Cell dimensions |  |  |  |  |
| *a*, *b*, *c* (Å) | 129.2, 162.8,<br>140.8 | 129.5, 163.9,<br>141.0 | 129.0, 162.0,<br>140.8 | 130.6, 165.9,<br>142.7 |
| $\alpha$, $\beta$, $\gamma$ (°) | 90.0, 90.0, 90.0 | 90.0, 90.0, 90.0 | 90.0, 90.0, 90.0 | 90.0, 90.0, 90.0 |
| Wavelength | 0.979 | 0.979 | 0.979 | 1.771 |
| Resolution (Å) | 53.17-3.62 (3.73-3.62) | 30.00-3.80 (3.94-3.80) | 40.0-3.53 (3.62-3.53) | 20.00-4.05 (4.14-4.05) |
| *R*<sub>meas</sub> (%) | 6.2 (41.6) | 6.2 (49.0) | 5.8 (58.9) | 10.4 (59.8) |
| *I*/σ*I* | 9.0 (1.7) | 8.4 (1.9) | 13.1 (1.7) | 11.1 (2.5) |
| Completeness (%) | 92.6 (90.7) | 95.5 (95.7) | 98.7 (86.7) | 98.1 (98.1) |
| Redundancy | 2.6 (2.2) | 3.6 (3.0) | 5.5 (3.2) | 10.1 (6.9) |
| CC<sub>1/2</sub> (%) | 99.9 (21.9) | 99.9 (28.3) | 99.9 (16.0) | 99.9 (20.3) |
| **Refinement** |  |  |  |  |
| Resolution (Å) | 53.26-3.62 (3.75-3.62) | 28.96-3.80 (3.94-3.80) | 38.91-3.53 (3.66-3.53) |  |
| No. reflections | 16974 (1529) | 14768 (1328) | 18289 (1615) |  |
| *R*<sub>work</sub>/*R*<sub>free</sub> (%) | 25.6 (37.8) / 26.3<br>(37.0) | 24.6 (40.3) / 26.9<br>(43.7) | 27.3 (41.4) / 28.4<br>(42.3) |  |
| No. atoms | 7703 | 7680 | 7678 |  |
| Protein | 7599 | 7598 | 7598 |  |
| Ligand/ion | 104 | 81 | 79 |  |
| Water | 0 | 1 | 1 |  |
| B-factors |  |  |  |  |
| Protein | 157.7 | 206.4 | 196.9 |  |
| Ligand/ion | 151.9 | 206.6 | 187.3 |  |
| Water | N/A | 225.0 | 182.0 |  |
| R.m.s deviations |  |  |  |  |
| Bond lengths (Å) | 0.009 | 0.009 | 0.008 |  |
| Bond angles (°) | 1.08 | 1.01 | 0.96 |  |
| Ramachandran plot |  |  |  |  |
| Favored (%) | 95.5 | 95.2 | 95.0 |  |
| Allowed (%) | 4.5 | 4.8 | 5.0 |  |
| Disallowed (%) | 0 | 0 | 0 |  |

The structure of the ts2 SERT variant in complex with paroxetine revealed only minor perturbations in the pose of the inhibitor and the conformations of residues involved in ligand binding in comparison to the ts3 - paroxetine structure (Fig. 4a,b). We believe this is because the hydroxyl group of threonine 439 faces subsite B, as does the serine at this position, and thus both are in a position to participate in interactions with ligands in the central site. We speculate that the ts2 variant has a higher affinity for paroxetine because, at least in part, the hydroxyl of Ser439 is closer to a ligand benzodioxol oxygen (3.9 Å) in comparison to the hydroxyl of Thr439 (5.2 Å). Further studies, at higher resolution, will be required to more definitively define transporter-ligand interactions.

**Figure 4.**
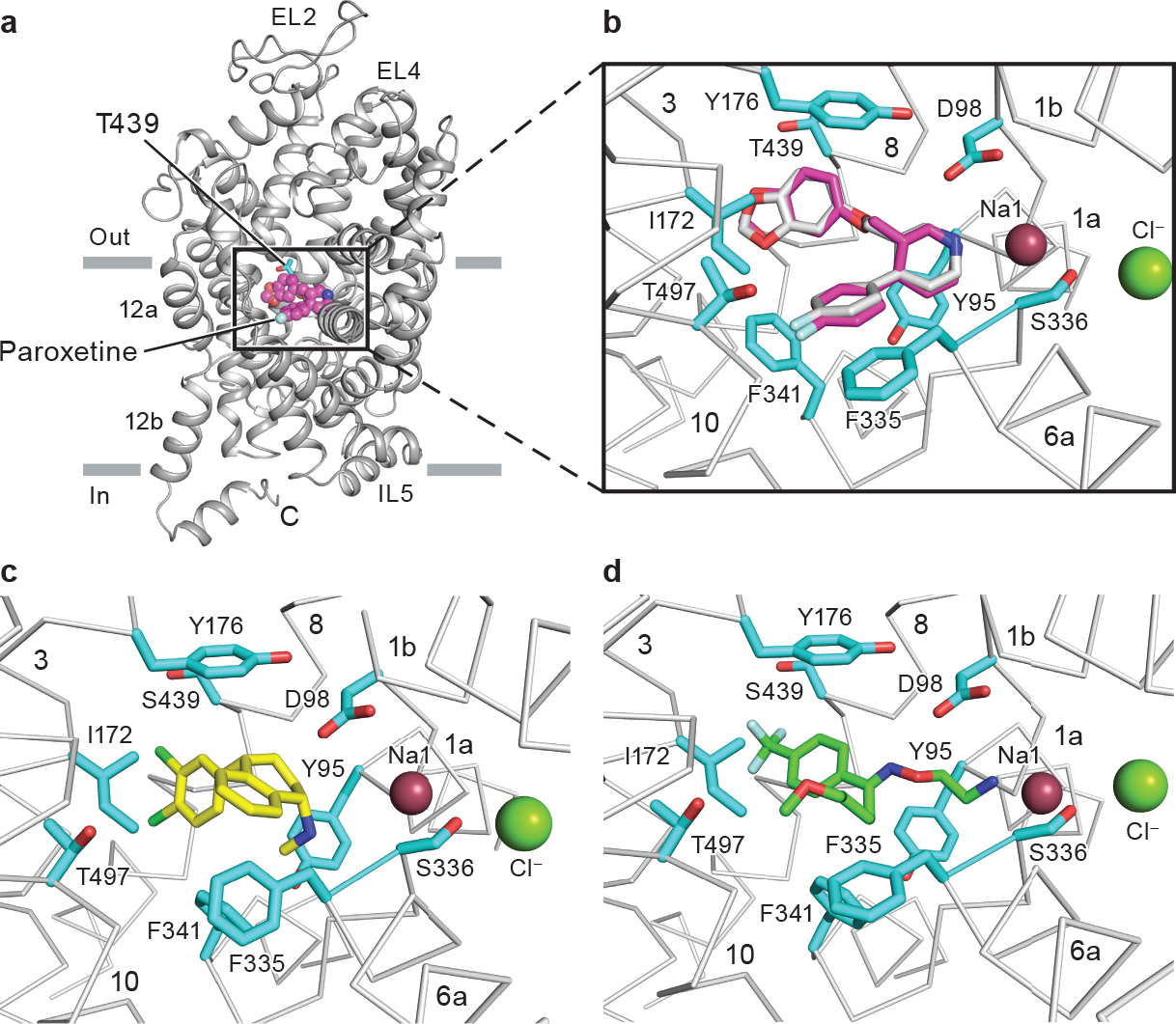
Antidepressant recognition. **a**, Overall view of the ts2 variant of the serotonin transporter in cartoon representation, Thr439 and paroxetine are depicted as sticks and spheres. **b**, Interactions of paroxetine (magenta) with residues in the central binding site of the ts2 variant. The position of paroxetine in the ts3 variant is overlaid in grey. **c**, Interactions of sertraline (yellow) with residues in the central binding site. **d**, Interactions of fluvoxamine (green) with residues in the central binding site.

The amine groups of sertraline and fluvoxamine also occupy subsite A, interacting with Asp98 and Try95 (Fig. 4c,d) residues crucial for drug binding^34–36^. In subsite B, Ser439 is within 4.5 and 3.9 Å of the halide atoms of sertraline and fluvoxamine while aromatic interactions are formed between Tyr176 and the drugs. The side chain of Ile172 adopts a similar conformation as it does in the paroxetine state, fitting snuggly between substituents in subsites B and C. In subsite C, Phe341 forms a face-to-face interaction with the naphthalene ring of sertraline while Phe335 forms a face-to-edge interaction with the naphthalene moiety. In the fluvoxamine complex, Phe341^25^ assumes a similar rotameric conformation as in the sertraline complex while the conformation of Phe335 is closer to that observed in the paroxetine structures.

Comparisons of the structures of the sertraline and fluvoxamine complexes to the previously determined structures of SERT with paroxetine and S-citalopram reveal important differences in the position of residues in subsite C. In the sertraline complex, relative to the paroxetine and S-citalopram complexes, Phe341 has flipped ‘upward’ to fill a space that is unoccupied by ligand and to interact with the naphthalene ring of sertraline; Phe335 undergoes a rotation of ~90° about its Cβ-Cγ bond (chi2) in order to accommodate the conformational change of Phe341 (Fig. 5a). In the fluvoxamine complex, Phe341 is in a similar position as the sertraline complex, but Phe335 adopts a conformation more similar to that observed in the paroxetine and S-citalopram complexes (Fig. 5b,c). We also note that Thr497 is also shifted by ~1 Å (Cα-Cα distance) in the sertraline and fluvoxamine complexes in comparison to the paroxetine complexes, likely because paroxetine is the only drug with a fluorinated substituent in subsite C, a substituent which interacts with Thr497.

**Figure 5.**
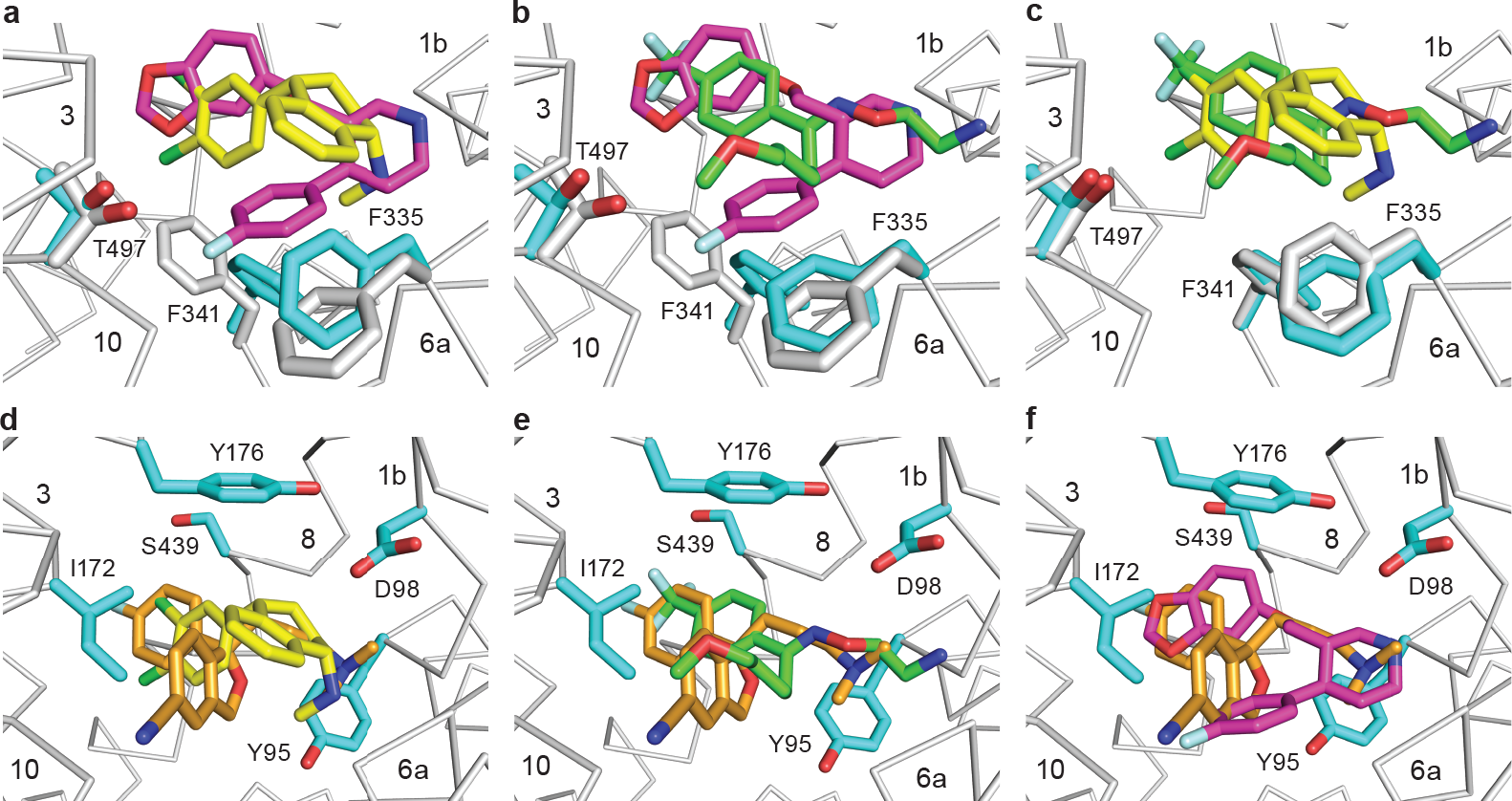
Comparisons of SSRI binding poses and central binding site structures. **a** Comparison of ts3 sertraline (yellow) and ts2 paroxetine (magenta) complexes. **b**, Comparison of ts3 fluvoxamine (green) and ts2 paroxetine binding. **c**, ts3 fluvoxamine *vs.* ts3 sertraline. **d**, Superposition of ts3 sertraline and ts3 *S*-citalopram (orange). **e**, ts3 fluvoxamine *vs.* ts3 S-citalopram. **f**, ts2 paroxetine *vs.* ts3 *S*-citalopram.

Superposition of all four SSRI complexes revealed different placement of the various drug substituents within the central binding site (Fig. 5d-f, Fig. 6) resulting in differences in van der Waals (>4 Å), aromatic (4.5-7 Å)^37^, and ionic (~4 Å)^38^ interactions. In subsite A, the distance from the amine of sertraline and S-citalopram to Asp98 is longer while the distance to the amine of fluvoxamine and paroxetine is shorter. Conversely, the positioning of the amine to Tyr95 is farther for fluvoxamine and paroxetine whereas the amine of *S*-citalopram and sertraline is more equally posed between Asp98 and Tyr95. In subsite B, the *meta* chlorine of sertraline and the fluorine of S-citalopram engages with Ser439 while the fluorines of fluvoxamine and the oxygen of paroxetine form even closer interactions with Ser439/Thr439 though the distance is not close enough for a hydrogen-bond to the halides^39^. The shorter distance to Ser439/Thr439 in the fluvoxamine complex and in the ts2 versus the ts3 paroxetine complexes may also explain, at least in part, the differences in affinity observed for fluvoxamine and fluoxetine in comparison to other SSRIs which do not have a trifluoro group in subsite B. The aromatic substituents of the SSRIs are arranged ~90° relative to Try176 and Ile172 in subsite B, but the distances to Tyr176 and Ile172 are divergent between inhibitors. In subsite C, the naphthalene group of sertraline is localized ‘higher’ than the benzofuran and fluorophenyl groups of *S*-citalopram and paroxetine.

**Figure 6.**
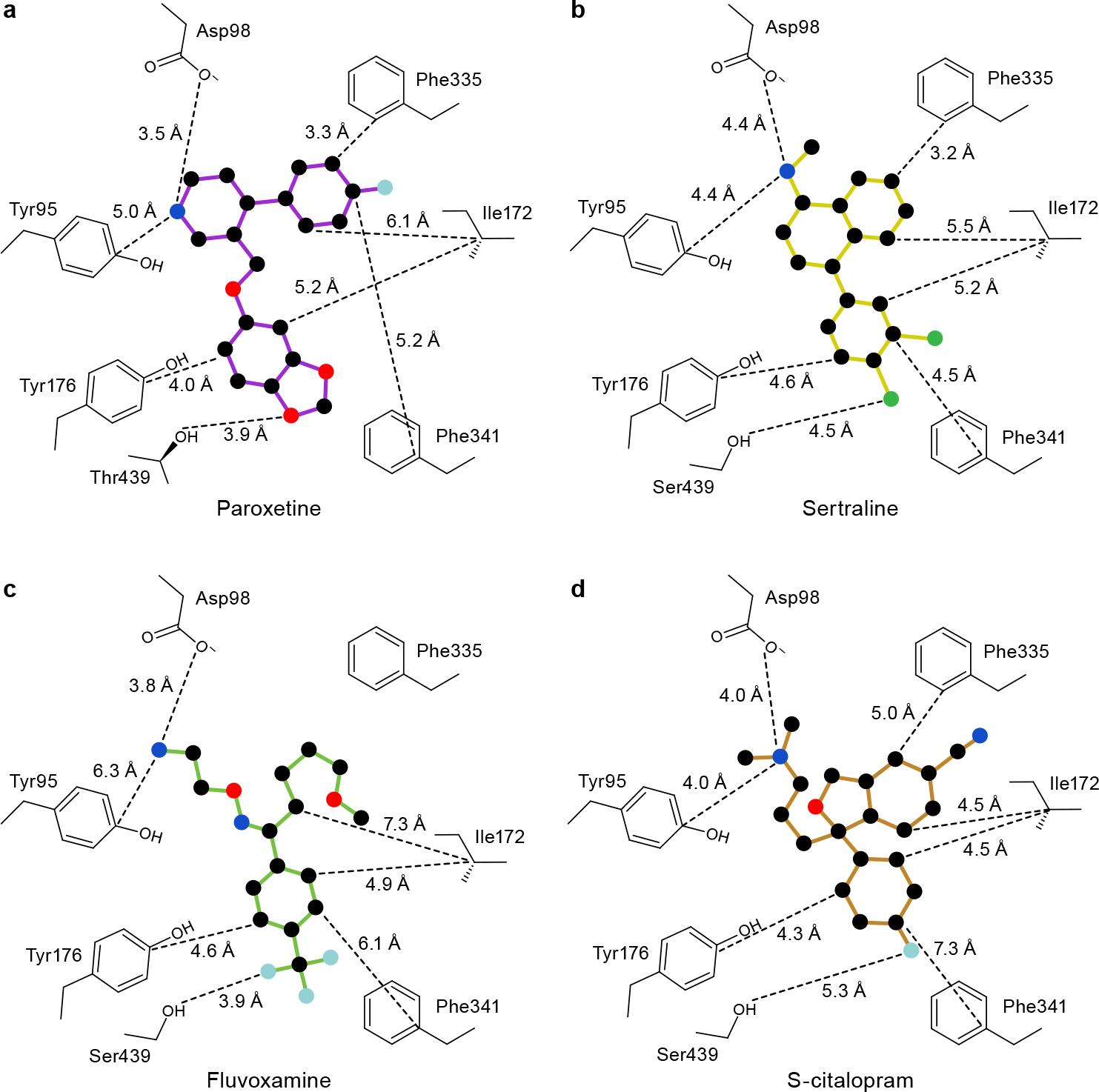
2D representation of selected drug binding residues. Distances of each residue to various positions of drug are shown for **a**, ts2 paroxetine **b**, ts3 sertraline **c**, ts3 fluvoxamine and **c**, ts3 *S*-citalopram.

Taken together, the SERT-SSRI complexes provide new insights into how pharmacophores of different drugs participate in interactions within the central binding site – all acting to stabilize the outward-open conformation of the transporter. We anticipate that these studies will provide a blueprint for the development of new therapeutic agents for the treatment of depression and anxiety disorders.

## STRUCTURE DEPOSITION

The coordinates for the structure have been deposited in the Protein Data Bank under the accession codes 6AWN (ts2-paroxetine), 6AWP (ts3-fluvoxamine), 6AWO (ts3-sertraline), 6AWQ (ts3-sertraline-anomalous) (see Table 1 for details).

## METHODS

### SERT constructs

The ts3 variant^19,20^ contains the thermostabilizing mutations Tyr110Ala, Ile291Ala, Thr439Ser, as well as mutation of surface-exposed cysteines Cys554, Cys580, and Cys622 to alanine. The ts3 SERT gene is then fused to sequences to express a C-terminal GFP fluorophore followed by a twin Strep and, lastly, a His_10_ purification tag. The mutation of Ser439 to threonine reverts the thermostabilizing mutation at position 439 of the ts3 construct to a threonine to then yield the ts2 construct.

### Expression and purification

SERT constructs were expressed by using baculovirus-mediated transduction of mammalian HEK293S GnTI^−^ cells^40^. Cells were solubilized in a buffer composed of 50 mM Tris (pH 8), 150 mM NaCl, 20 mM *n*-dodecyl-β-D-maltoside (DDM), and 2.5 mM cholesterol hemisuccinate (CHS). Detergent-solubilized SERT was purified via Strep-Tactin affinity chromatography in a buffer composed of 20 mM Tris pH 8, 100 mM NaCl (TBS) containing 1 mM DDM, 0.2 mM CHS, 5% glycerol, 25 μM lipid (1-palmitoyl-2-oleoyl-sn-glycero-3-phosphocholine, 1-palmitoyl-2-oleoyl-sn-glycero-3-phosphoethanolamine, and 1-palmitoyl-2-oleoyl-sn-glycero-3-phosphoglycerol at a molar ratio of 1:1:1). Following elution from the column, the resulting protein was digested by thrombin and EndoH. SERT was then combined with the recombinant antibody fragment (Fab) 8B6 in a molar ratio of 1:1.2 (SERT:8B6) and the complex was purified by size-exclusion chromatography in TBS supplemented with 40 mM n-octyl β-D-maltoside, 0.5 mM CHS, 5% glycerol and 25 μM of the lipid mixture described above. Prior to crystallization, the purified SERT-8B6 complex was concentrated to 2 mg/ml; inhibitors were added to the protein solution
to reach a final concentration of 50 μM and an additional amount of 8B6 Fab was added so that the final concentration of the added Fab was 1 μM.

### Crystallization

The SERT-Fab complex crystals were grown by hanging drop vapor diffusion at 4 °C using a ratio of 2 μl protein to 1 μl reservoir solution, the latter of which contained 50-100 mM Tris pH 8.5, 25-75 mM Li_2_SO_4_, 25-75 mM Na_2_SO_4_, 33.5 – 36 % PEG 400 and 0.5% 6-aminohexanoic acid.

### Data collection and structure refinement

Crystals were directly flash cooled in liquid nitrogen. Data was collected at the Advanced Photon Source (Argonne National Laboratory, beamline 24-ID-E) and at the Advanced Light Source (Lawrence Berkeley National Laboratory, beamline 5.0.2). X-ray data sets were processed using XDS^41^. Molecular replacement was performed using coordinates (PDB 5I6X code) from the prior structure determination of SERT^8^ using the computer program PHASER^42^. Several iterations of refinement and manual model building were carried out using PHENIX^43^ and Coot^44^ until the models converged to acceptable R-factors and stereochemistry. Polder ‘omit’ maps were calculated by excluding bulk solvent as previously described^33^.

### Radioligand binding assays

Saturation binding experiments employed the scintillation proximity assay (SPA)^45^ and a solution containing 10 nM SERT, 0.5 mg/ml Cu-Ysi beads, TBS, 1 mM DDM, 0.2 mM CHS, and [^3^H-paroxetine] from 0.15-20 nM. Competition binding experiments were also performed using SPA, in the same buffer as the saturation binding experiments and ligands at concentrations of 5 nM for ^3^H-paroxetine and at 0.01-100,000 nM for the cold competitors. Each data point was measured in triplicate and each experiment was performed three times. The error bars for each data point represent the standard error of the mean. K_i_ values were determined using the Cheng-Prusoff equation^46^ in Graphpad Prism.

## ACKNOWLEDGEMENTS

We thank L. Vaskalis for assistance with figures, H. Owen for help with manuscript preparation and all Gouaux laboratory members for discussion. We acknowledge the staff of the Berkeley Center for Structural Biology at the Advanced Light Source and the Northeastern Collaborative Access Team at the Advanced Photon Source for assistance with data collection. J.A.C. has support from a Banting postdoctoral fellowship from the Canadian Institutes of Health Research. We are particularly grateful to Bernie and Jennifer LaCroute for their generous support, as well as for funding from the National Institutes of Health (NIH) (5R37MH070039). E.G. is an Investigator with the Howard Hughes Medical Institute.

## Author contributions

J.A.C. and E.G. designed the project. J.A.C. performed protein purification, crystallography, and biochemical assays. J.A.C. and E.G. wrote the manuscript.

The authors declare no competing financial interests.

---

**Extended Data Figure 1.**
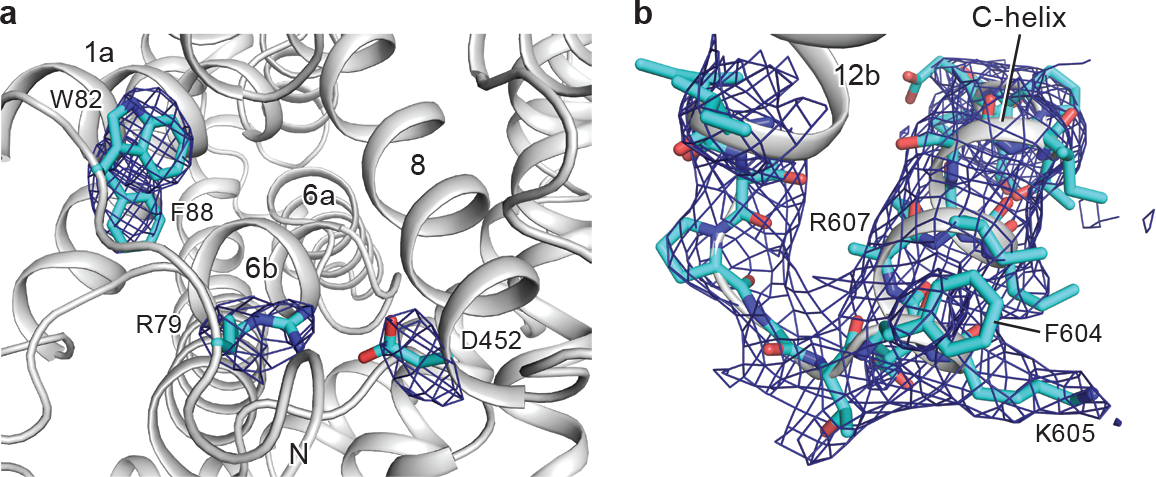
Intracellular gate and C-terminus. **a**, Positions of residues associated with the intracellular gate are shown and the Polder ‘omit’ electron density is shown in blue mesh (contoured at 6-8σ). **b**, Fit of C-terminal residues into 2F_o_ − F_c_ electron density map (blue mesh), contoured at 0.75σ. Density for most side chains of the SEC24C recognition sequence were observed except for Arg607.

**Extended Data Figure 2.**
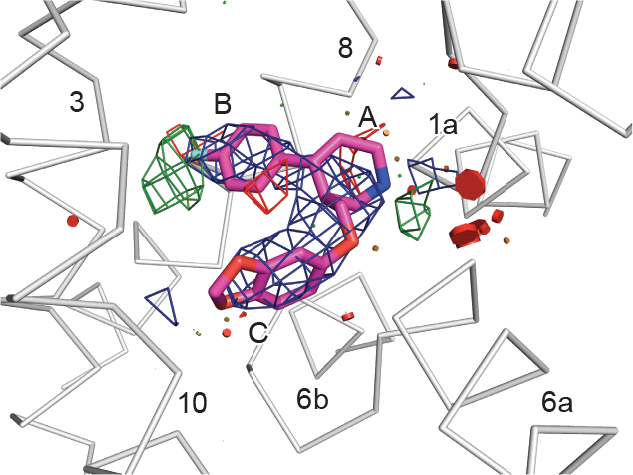
Fitting of paroxetine with fluorophenyl in subsite B and benzodioxol in subsite C. Shown is the 2F_o_-F_c_ (blue mesh) and F_o_-F_c_ (positive, green; negative red) electron density associated with paroxetine after refinement, contoured at 2σ and 2.5σ respectively. Shown in red to yellow to green disks are severe to significant to slight overlaps of van der Waals radii of residues within 5 Å of paroxetine. The size of the disk also indicates the degree of clashing.

**Extended Data Figure 3.**
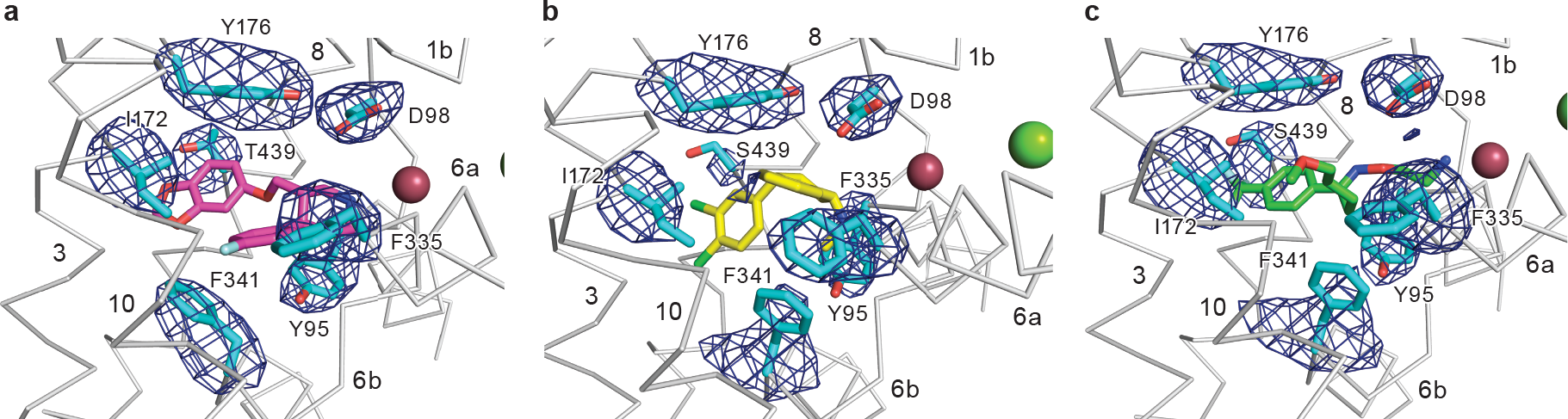
Electron density of drug binding residues. **a**, Residues associated with paroxetine (magenta) binding are shown and the Polder ‘omit’ electron density is shown in blue mesh (contoured at 7σ). **b**, Residues associated with sertraline (yellow) binding are shown and the Polder ‘omit’ electron density is shown in blue mesh (contoured at 6-7σ). **c**, Residues associated with fluvoxamine (green) binding are shown and the Polder ‘omit’ electron density is shown in blue mesh (contoured at 5-6σ).

